# Functional instability allows access to DNA in longer Transcription Activator-Like Effector (TALE) arrays

**DOI:** 10.1101/319749

**Authors:** Kathryn Geiger-Schuller, Jaba Mitra, Taekjip Ha, Doug Barrick

**Affiliations:** T.C. Jenkins Department of Biophysics and Program in Molecular Biophysics, Johns Hopkins University, 3400 N. Charles St, Baltimore MD 21218.; Current address: Broad Institute of Harvard and Massachusetts Institute of Technology, Cambridge, MA 02142.; Materials Science and Engineering, University of Illinois Urbana-Champaign, Urbana IL 61801.; Department of Physics, Center for the Physics of Living Cells and Institute for Genomic Biology, University of Illinois at Urbana Champaign, Urbana IL 61801.; Department of Biophysics and Biophysical Chemistry and Department of Biomedical Engineering, Johns Hopkins University, Baltimore MD 21205.; Howard Hughes Medical Institute, Baltimore, MD, 21205, USA.

**Keywords:** TALE repeat, Single-molecule biophysics, FRET, Functional instability, Deterministic modeling

## Abstract

Transcription activator-like effectors (TALEs) bind DNA through an array of tandem 34-residue repeats. Here, we examine the kinetics of DNA binding for a set of TALE arrays with varying numbers of identical repeats using single molecule microscopy. Using a new deterministic modeling approach, we find evidence for conformational heterogeneity in both the free- and DNA-bound TALE arrays. Combined with previous work demonstrating populations of partly folded TALE states, our findings reveal a functional instability in TALE-DNA binding. For TALEs forming less than one superhelical turn around DNA, partly folded open states inhibit DNA binding. In contrast, for TALEs forming more than one turn, the partly folded open states facilitate DNA binding. Overall, we find that increasing repeat number results in significantly slower interconversion between the various DNA-free and DNA-bound states. These findings highlight the role of conformational heterogeneity and dynamics in facilitating macromolecular complex assembly.

**Impact Statement:** Single molecule DNA-binding trajectories and deterministic modeling analyses demonstrate a functional role for high energy partly folded states in Transcription Activator-Like Effectors (TALEs) that could improve future TALEN design.

## Introduction

Transcription activator-like effectors (TALEs) are bacterial proteins containing a domain of tandem DNA-binding repeats as well as a eukaryotic transcriptional activation domain (Kay et al., 2007; Römer et al., 2007). The repeat domain binds double stranded DNA with a register of one repeat per base pair. Specificity is determined by the sequence identity at positions twelve and thirteen in each TALE repeat, which are referred to as repeat variable diresidues (RVDs) (Boch et al., 2009; Miller et al., 2015; Moscou and Bogdanove, 2009). This specificity code has enabled design of TALE-based tools for transcriptional control (Cong et al., 2012; Geissler et al., 2011; Li et al., 2012; Mahfouz et al., 2012; Zhang et al., 2011), DNA modifications (Maeder et al., 2013), in-cell microscopy (Ma et al., 2013; Miyanari et al., 2013), and genome editing (TALENs) (Christian et al., 2010; Li et al., 2011).

TALE repeat domains wrap around DNA in a continuous superhelix of 11.5 TALE repeats per turn (Deng et al., 2012; Mak et al., 2012). Because TALEs contain on average 17.5 repeats (Boch and Bonas, 2010), most form over 1.5 full turns around DNA. Many multisubunit proteins that form rings around DNA require energy in the form of ATP to open or close around DNA (reviewed in O’Donnell and Kuriyan, 2006), yet TALEs are capable of wrapping around DNA without energy from nucleotide triphosphate hydrolysis. One possibility is that TALEs bind DNA through an energetically accessible open conformation. Consistent with this possibility, we previously demonstrated that TALE arrays can populate partly folded or broken states (Geiger-Schuller and Barrick, 2016). While the calculated populations of partly folded states in TALE repeat arrays are small, these populations are many orders of magnitude larger than calculated populations of partly folded states in other previously studied repeat arrays (consensus Ankyrin (Aksel et al., 2011) and DHR proteins (Geiger-Schuller et al., 2018)) suggesting a potential functional role for the high populations of partially folded states in TALE repeat arrays.

Consensus TALEs (cTALEs) are homopolymeric arrays composed of the most commonly observed residue at each of the 34 positions of the repeat (Geiger-Schuller and Barrick, 2016). In addition to simplifying analysis of folding and conformational heterogeneity in this study, the consensus approach simplifies analysis of DNA binding, eliminating contributions from sequence heterogeneity and providing an easy means of site-specific labeling.

Here we characterize DNA binding kinetics of cTALEs using total internal reflection fluorescence single-molecule microscopy. We find that consensus TALE arrays bind to DNA reversibly, with high affinity. Analysis of the dwell-times of the on- and off-states reveals multiphasic binding and unbinding kinetics, suggesting conformational heterogeneity in both the free and DNA bound state. We develop a deterministic optimization analysis that supports such a model, and provides rate constants for both conformational changes and binding. Comparing the dynamics observed here to previously characterized local unfolding suggests that locally unfolded states inhibit binding of short cTALE arrays (less than one full superhelical turn around DNA), whereas they promote binding of long arrays (more than 1 full superhelical turn). Whereas local folding of transcription factors upon DNA binding is well documented (Spolar and Record, 1994), local unfolding in the binding process has not. Our results present a new mode of transcription factor binding where the major conformer in the unbound state is fully folded, requiring partial unfolding prior to binding. The critical role of such high energy partly folded states is an exciting example of functional instability.

## Results

### cTALE design

Consensus TALE (cTALE) repeat sequence was design previously described (Geiger-Schuller and Barrick, 2016). To avoid self-association of cTALE arrays, we fused arrays to a conserved N-terminal extension of the PthXo1 gene. Although the sequence of this domain differs from a TALE repeat sequence, the structure of this domain closely mimics four TALE repeats (Gao et al., 2012; Mak et al., 2012) and is required for full transcriptional activation (Gao et al., 2012). In this study, all repeat arrays contain this solubilizing N-terminal domain.

### cTALE local instability promotes population of partly folded states

Figure 1A depicts types of partly folded states of a generic repeat protein. In the fully folded state, all repeats are folded, and all interfaces are intact. In the end-frayed states, one or more terminal repeats are unfolded and all interfaces, except the interface(s) between the unfolded and adjacent folded repeat(s), are intact. In the internally unfolded states, a central repeat is unfolded and all interfaces, except the interfaces involving the unfolded repeats, are intact. In the interfacially ruptured state, all repeats are folded but one interface is disrupted due to local structural distortion.

**Figure 1.**
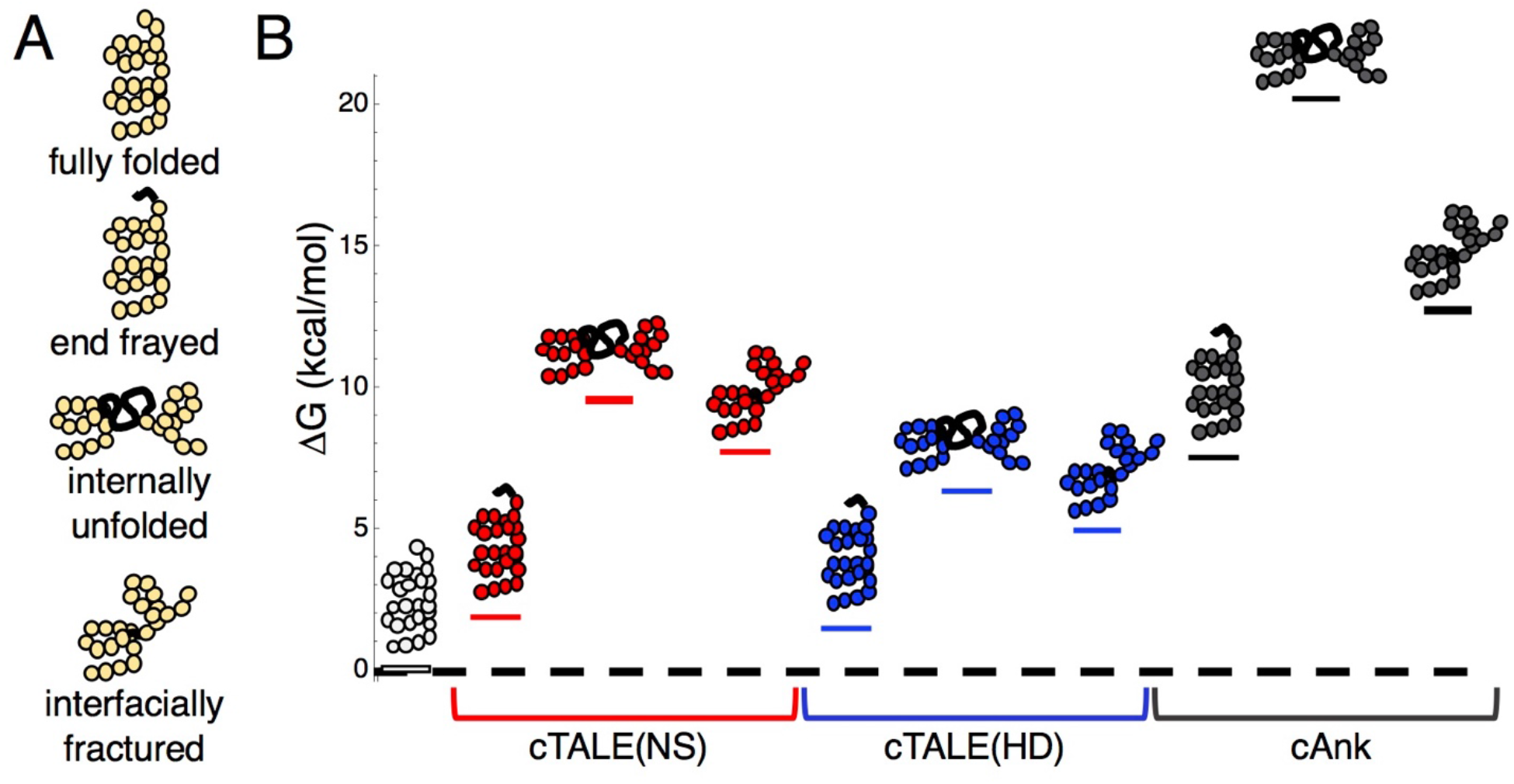
cTALEs populate partly folded states. (A) Cartoon of different partly folded TALE conformational states. End-frayed states have one (or more) terminal repeats unfolded. Internally unfolded states have a central repeat unfolded. Interfacially fractured states have a disrupted interface between adjacent repeats. (B) Free energies of partly folded states, calculated from previously published measurements and analysis (Geiger-Schuller and Barrick, 2016), relative to the fully folded state, for consensus TALE repeats with the NS repeat-variable diresidue sequence (cTALE(NS), red), consensus TALE repeats with the HD repeat-variable diresidue sequence (cTALE(HD), blue), and consensus ankyrin repeats (cAnk, black).

Figure 1B shows calculated free energy differences between various partly folded states and the fully folded repeat array for two different RVDs (NS and HD) in an otherwise identical consensus sequence background, using the intrinsic and interfacial engeries we determined previously (Geiger-Schuller and Barrick, 2016). The distribution of partly folded states is calculated for different 20-repeat arrays containing two types of TALE arrays (with the NS RVD in red and with the HD RVD in blue) as well as consensus ankyrin arrays (cAnk in black). For cTALE arrays, end frayed states are within a few *RT* of the folded state, internally unfolded states are highest in energy, and interfacially ruptured states fall energetically between end frayed and internally unfolded states.

Changing the RVD sequence affects the distribution of these partly folded states: HD repeat containing arrays are more likely to internally unfold or interfacially rupture than NS repeat containing arrays. However, both types of cTALEs are more likely to populate many of these partly folded states than cAnk is to populate even the lowest energy partly folded state, the end frayed state. Thus, compared to ankyrin repeats, cTALEs are locally unstable, meaning they are likely to form partly folded states. As these states disrupt the superhelix, they may facilitate DNA binding.

### Single-molecule studies of cTALE binding to DNA

To ask if cTALE local instability is relevant for DNA binding kinetics, DNA binding trajectories were measured using single molecule total internal reflection fluorescence (smTIRF). Figure 2A shows a schematic of the smTIRF experiments performed to measure DNA binding. For site-specific cTALE labeling, R30 is mutated to cysteine in a single repeat. Position 30 is frequently a cysteine in naturally occurring TALEs (in earlier folding studies, arginine was chosen in the consensus sequence to avoid disulfide formation) (Geiger-Schuller and Barrick, 2016). This cysteine was Cy3 (FRET donor)-labelled using maleimide chemistry, and was attached to biotinylated slides via the C-terminal His_6_ tag and α-Penta•His antibodies. At salt concentrations below 300 mM NaCl, cTALEs aggregate. Because DNA binding is weak at high salt concentrations, measuring binding kinetics in bulk at high salt is not possible. However, tethering cTALEs to the quartz slide at high salt prevents self-association, even at the low salt concentrations required to study DNA binding kinetics. A histogram of NcTALE _8_ (8 NS-type repeats and the N-terminal domain) labeled via a cysteine in the first repeat shows a single peak at zero FRET efficiency, as expected for donor-only constructs (Figure 2B).

**Figure 2.**
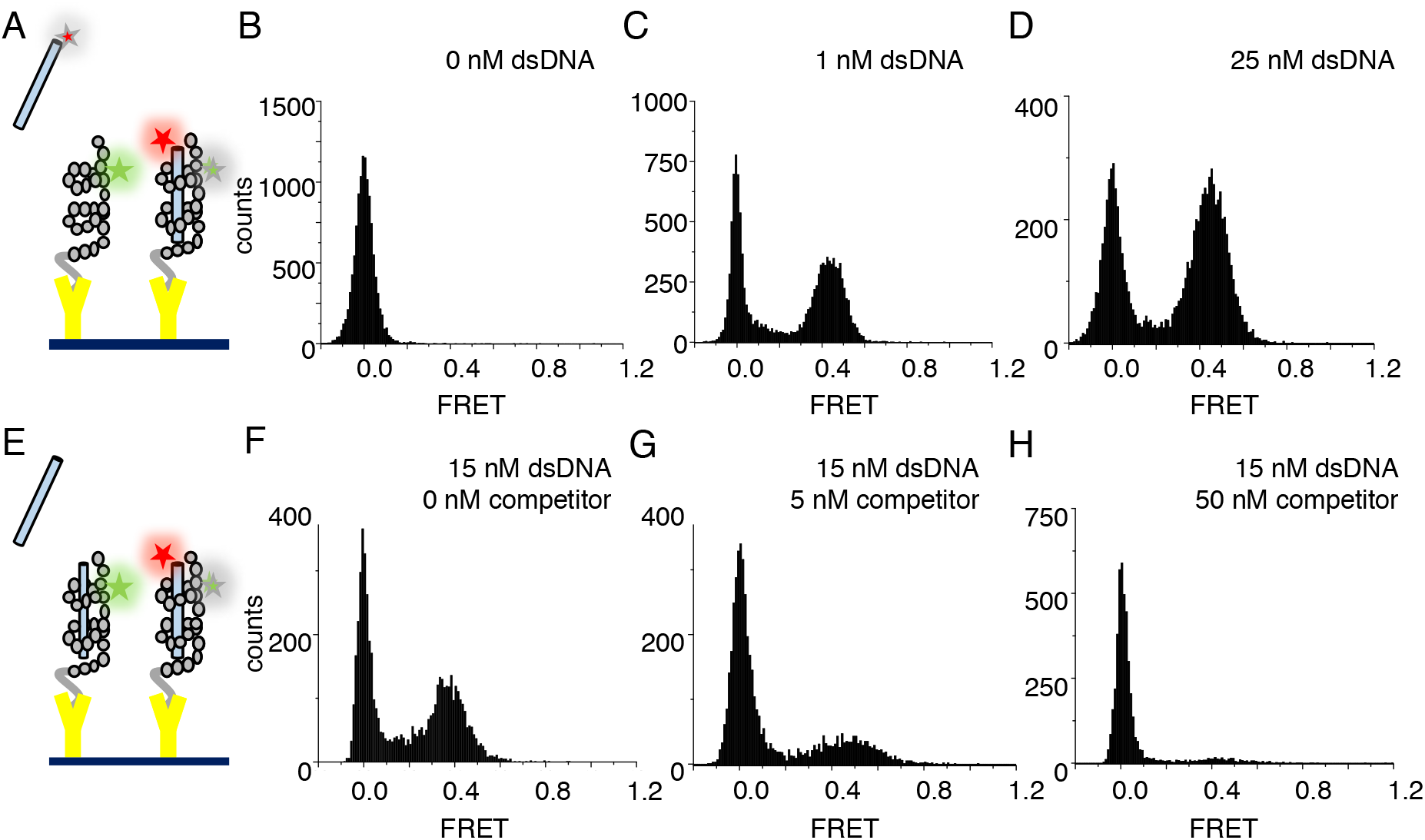
cTALEs bind dsDNA reversibly. (A) Schematic of single-molecule FRET assay, with donor-labelled cTALE attached to a surface, and acceptor-labelled DNA free in solution. (B-D) Single molecule FRET histograms show the appearance of a peak at a FRET efficiency of 0.45 with increasing labelled DNA, consistent with a DNA-bound cTALE. (E) Schematic of single-molecule FRET competition assay, with donor-labelled cTALE attached to a surface, and acceptor-labelled DNA as well as competitor unlabeled DNA free in solution. (F-H) Single molecule FRET histograms show the disappearance of the peak at 0.45 FRET efficiency with increasing unlabeled competitor DNA. Conditions: 20 mM Tris pH 8.0, 200 mM KCl.

To test for DNA binding to tethered cTALE constructs, we added Cy5 (FRET acceptor)-labeled 15 bp-long DNA (Cy5.A_15_ /T_15_) to tethered NcTALE_8_. This results in a new peak at a FRET efficiency of 0.45, indicating that DNA binds directly to cTALE arrays. As DNA concentration in solution is increased, the peak at 0.45 FRET efficiency increases in population (Figure 2C-D), suggesting a measurable equilibrium between free and bound DNA rather than saturation or irreversible binding. In support of this, single molecule time trajectories show interconversion between bound and unbound states, providing access to rates of binding and dissociation.

As expected for reversible complex formation, the peak at 0.45 FRET efficiency can be competed away by adding mixtures of labeled and unlabeled DNA to pre-formed cTALE-labelled DNA complexes (schematic shown in Figure 2E; pre-formed complex shown in Figure 2F, competition data shown in Figure 2G-H). Challenging pre-formed complexes with a mixture of 5 nM unlabeled DNA and 15 nM labeled DNA results in a slight decrease in the population of the peak at 0.45 FRET (compare Figures 2F and 2G). Challenging with a mixture of 50 nM unlabeled DNA and 15 nM labeled DNA further decreases the peak at 0.45 FRET (Figure 2H).

### cTALE arrays display multiphasic DNA-binding kinetics

In addition to the short smTIRF movies used to generate smFRET histograms from many molecules, long movies were also collected to examine the extended transitions of individual molecules between the low- and high-FRET (0 and 0.45) (Figure 3A-B). A transition from low to high FRET (0 to 0.45) indicates that the acceptor fluorophore on DNA moved close enough to the donor on the protein for FRET and is likely a binding event. A transition from high to low FRET (0.45 to 0.0) indicates the acceptor fluorophore on DNA moved too far away from the donor on the protein for FRET and is likely an unbinding event. Low-FRET states show low colocalization with signal upon direct excitation of the acceptor, confirming that high-FRET states are DNA-bound states and low-FRET states are DNA-free states (Figure 3-figure supplement 1). These long single molecule traces show both long- and short-lived low- and high-FRET states, indicating that kinetics are multi-phasic (Figure 3A-B). Binding events (transitions from low to high FRET) become more frequent as bulk DNA concentration increases (compare representative traces at 1 nM dsDNA to 15 nM dsDNA; Figure 3A and Figure 3B). Cumulative distributions generated from dwell times in the low FRET state at a given DNA concentration are best-fit by a double-exponential decay, indicating a minimum of two kinetic phases associated with binding events (Figure 3C). Cumulative distributions generated from dwell times in the high FRET state are best-fit by a double-exponential decay, indicating that there are a minimum of two kinetic phases for unbinding as well (Figure 3D).

**Figure 3.**
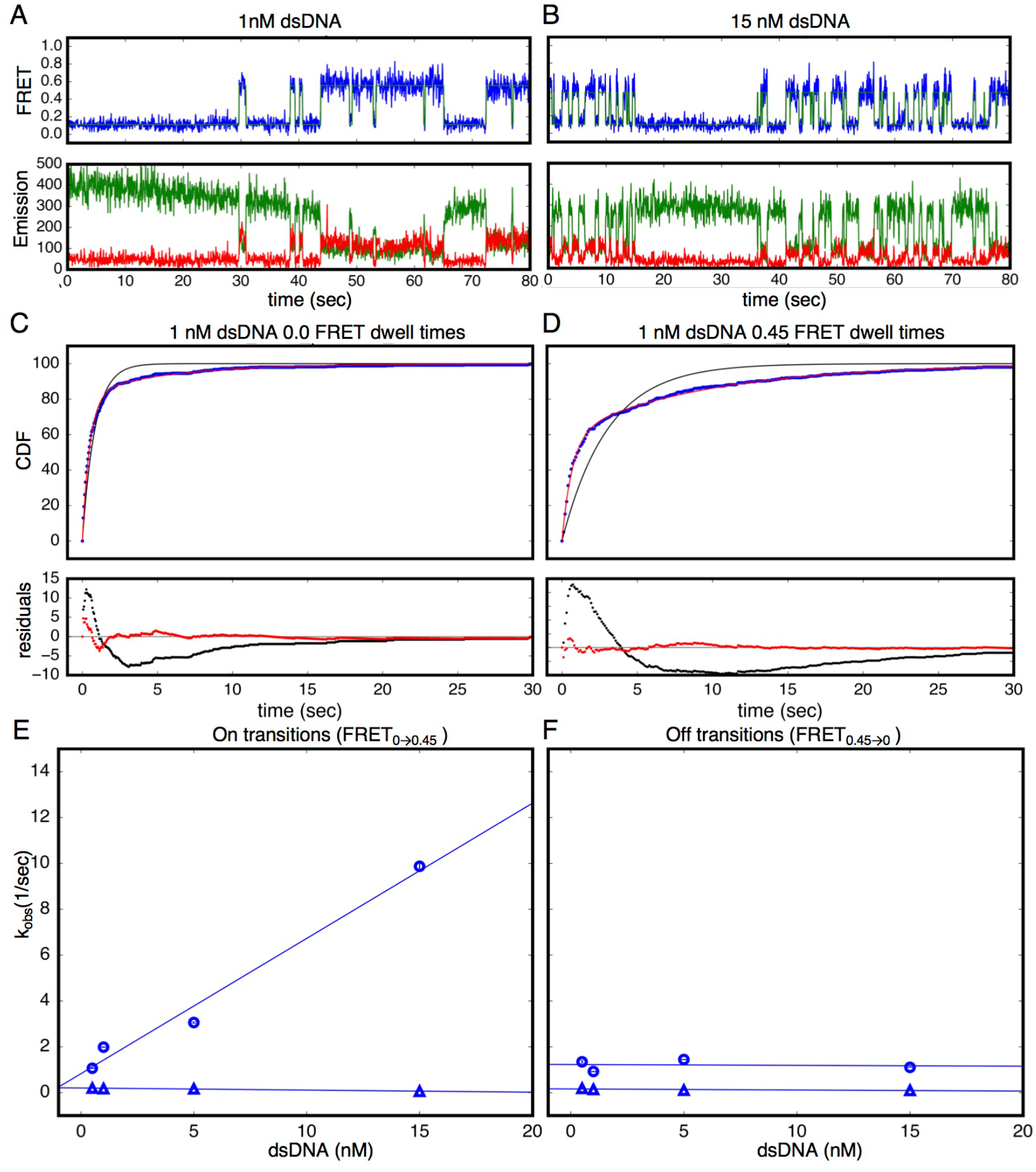
Single Molecule (SM) kinetics show multiple phases in binding and unbinding kinetics. (A-B) Long time trajectories showing transitions between low- and high-FRET states (efficiencies of 0 and 0.45). The top panel shows calculated FRET efficiency in blue and two-state Hidden Markov Model fit in green (McKinney et al., 2006). The bottom panels show Cy3 and Cy5 fluorescence emission in green and red respectively. At low DNA concentration (A), the low FRET state predominates. As DNA concentration is increased (B), more time is spent in the high FRET state, because the dwell times in the low FRET state are shorter. At low DNA concentrations, there appears to be long- and short-lived high-FRET states. Likewise, at near-saturating DNA concentrations, there appear to be long and short-lived low FRET states. (C, D) Cumulative distributions of low- and high-FRET dwell times (blue circles). Fits to single-exponentials (black) show large nonrandom residuals (lower panels), consistent with the heterogeneity noted in (A) and (B). Double-exponentials (red) give smaller, more uniform residuals. (E) Apparent association rate constants as a function of DNA concentration. The apparent rate constants for the fast phase depend on DNA concentration (blue circles), indicating a bimolecular step binding event. The apparent rate constants for the slow phase do not depend on DNA-concentration (blue triangles), suggesting an isomerization event. (F) Apparent dissociation rate constants as a function of DNA concentration (phase 1 shown in blue circles, and phase 2 shown in blue triangles). Neither phase shows a DNA concentration dependence, indicating a dissociation and/or isomerization events. 67.4% confidence intervals are estimated using the conf_interval function of lmfit by performing F-tests (Newville et al., 2014). Conditions: 20 mM Tris pH 8.0, 200 mM KCl.**Figure supplement 1**. Alternating laser experiments show agreement between cTALE_8_ FRET and colocalization kinetics. **Figure supplement 2**. cTALEs do not slide onto ends of short dsDNA. **Source data 1.** List of values used to construct long time trajectories. **Source data 2.** List of values used to construct CDFs.

The rate constant for the fast phase in DNA binding shows a linear increase with DNA concentration (Figure 3E), indicating that this step involves an associative binding mechanism. The slope of these rate constants as a function of DNA concentration gives a bimolecular rate constant of 5.9×10^8^nM^−1^s^−1^, close to the diffusion limit. The rate constant for the slower phase (0.59 s^−1^) is independent of DNA concentration indicating a unimolecular isomerization mechanism (Figure 3E).

In contrast, neither of the two fitted rate constants for transitions from high to low FRET (0.45 to 0.0; unbinding events) depends on DNA concentration, suggesting that unbinding involves two (or more) unimolecular processes (Figure 3F). The rate constants of these two phases are 1.2 s^−1^and 0.13 s^−1^respectively.

To rule out kinetic contributions of TALEs threading axially onto the ends of short DNAs, binding kinetics were measured with capped double-helical DNA sites. Capped DNA was generated by forming 5’digoxygenin-A_5_-Cy5-A _15_ duplexed with 5’-digoxygenin-T_26_ and adding a three-fold molar excess of anti-Digoxygenin. Low and high FRET dwell time cumulative distributions generated from capped DNA-binding kinetics are bi-phasic, similar to distributions from uncapped DNA (Figure 3-figure supplement 2). The DNA concentration-independent rate constant for binding is roughly the same for capped DNA as for uncapped DNA (compare FRET _L→H_ red and blue triangles in Figure 3 figure supplement 2), as are the dissociation rate constants (compare FRET _H→L_ red and blue triangles in as well as FRET _H→L_ red and blue circles in Figure 3-figure supplement 2). The rate constant for bimolecular binding of capped DNA decreases compared to that for uncapped DNA (compare FRET _L→H_ red and blue circles in Figure 3-figure supplement 2), which is consistent with the expected decrease in the rate of diffusion of the larger capped DNA. To assess the effect of molecular weight increase on diffusion of capped versus uncapped DNA, Sednterp (edited by S. E. Harding, 1992), a program commonly used to estimate sedimentation and diffusion properties of biomolecules, was used to estimate maximum diffusion coefficients. Including the two antibodies bound on the ends of capped DNA (320 kDa total) gives an estimated diffusion coefficient of 4.7×10^−7^cm^2^s^−1^, which is much lower than the estimated diffusion coefficient of the uncapped DNA (1.5 x10^−6^cm^2^s^−1^). This ~3.6-fold decrease in the diffusion constant for capped DNA is similar to the 6.7-fold decrease in the bimolecular rate constant for binding of capped DNA (Figure 3-figure supplement 2).

### Longer cTALEs have slower DNA binding and unbinding kinetics

To examine how increasing the length of the cTALE array influences DNA binding, we generated Cy3-labelled constructs with 16 and 12 cTALE repeats, and measured binding to a longer Cy5-labelled DNA (A_23_/T_23_). Because we did not observe FRET for these longer constructs, a fluorescence colocalization microscopy protocol was used instead (Figure 4-figure supplement 1). In this protocol, Cy3 was first imaged for ten camera frames (1017.5 msec total) to identify positions of single TALE molecules. Then a long time series of fluorescence images of Cy5 signal were collected through directly exciting Cy5 on the DNA, and time trajectories of Cy5 signal were generated from the initially identified single TALE positions.

**Figure 4.**
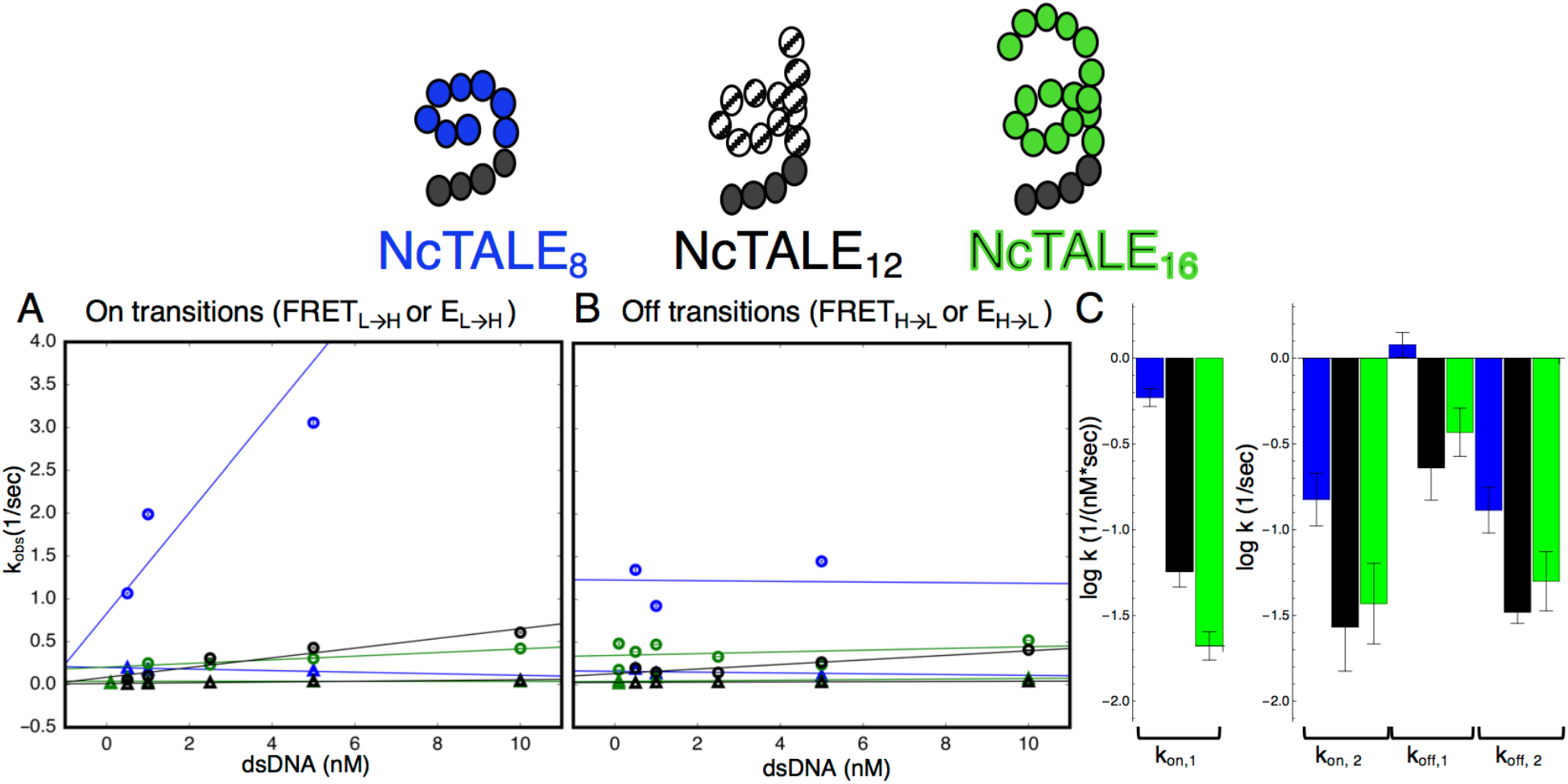
A 16-repeat TALE protein binds and unbinds DNA more slowly than an eight repeat protein. (A) Apparent association rate constants as a function of DNA concentration for an 8 repeat cTALE (blue), a 12 repeat cTALE (black), and a 16 repeat cTALE (green). 8 repeat TALE kinetics are measured by FRET (FRET_L→H_) while 12 and 16 repeat TALE kinetics are measured by colocalization (EL_→_ H). The apparent rate constants for the fast phase of binding are DNA concentration dependent (blue, black, and green circles), indicating a bimolecular binding event. The DNA concentration-dependence is greatest (larger slope) for the 8 repeat cTALE. The apparent rate constants for the slow phase do not depend on DNA-concentration (blue, black, and green triangles), suggesting an isomerization event. (B) Apparent dissociation rate constants as a function of DNA concentration (phase 1 shown in circles, and phase 2 shown in triangles). Neither phase shows a DNA concentration dependence, indicating a dissociation and/or isomerization events. Rate constants for all phases are slower for the 12-repeat construct (black) and 16-repeat construct (green) than for the 8-repeat construct (blue), particularly for the bimolecular binding step. (C) Log _10_ of rate constants for 8 (blue), 12 (black), and 16(green) repeat cTALEs. Units of the bimolecular binding rate constant are nM^−1^s^−1^, other unimolecular rate constants have units s^−1^. 67.4% confidence intervals are estimated using the conf_interval function of lmfit by performing F-tests (Newville et al., 2014). Conditions: 20 mM Tris pH 8.0, 200 mM KCl. **Figure supplement 1**. Colocalization trajectories show TALE-DNA binding and unbinding events. **Figure supplement 2**. Bound and unbound lifetimes of 16- and 12-repeat TALE proteins are consistent with multiphasic binding and unbinding. **Figure supplement 3**. Distance estimates between labeling sites for NcTALE _8_ and NcTALE _16_ and the 5’ ends of bound DNA.

Increasing the number of cTALE repeats from 8 to 12 and 16 dramatically affects DNA binding kinetics. Long movies collected over a range of DNA concentrations show short- and long-lived Cy5 signal on and off states, indicating a level of kinetic heterogeneity similar to NcTALE_8_ (Figure 4-figure supplement 1). Single molecule traces were analyzed using a thresholding filter (see Materials and Methods and Figure 4-figure supplement 1) to identify states and dwell times. Cumulative distributions were generated from dwell times at low Cy5 signal (unbound states, with lifetimes representing binding kinetics), and at high Cy5 signal (bound states, with dwell times representing unbinding kinetics). As with the eight repeat constructs, unbound cumulative distributions for these longer TALE arrays are best-fit by double exponential decays, particularly at high DNA concentrations (compare the cumulative distribution at low DNA concentration, Figure 4-figure supplement 2A, to cumulative distribution at 5 nM DNA, Figure 4-figure supplement 2B). Bound cumulative distributions for longer TALE arrays are best-fit by double exponential decays (Figure 4-figure supplement 2C-D). All apparent rate constants are much smaller for NcTALE_16_ and NcTALE_12_ (green/black circles and triangles, Figure 4A-B) compared to NcTALE_8_, indicating that binding and unbinding is impeded by increasing the length of the binding surface between cTALEs and their cognate DNA (Figure 4C). To address whether differences in binding kinetics are related to experimental differences between colocalization and FRET assays, alternating laser experiments were performed by switching between FRET and colocalization detection (every 5 frames) within single molecule trajectories (Figure 3-figure supplement 1). Changes in FRET and colocalization signals occurred simultaneously according to single molecule time traces, showing that differences in binding and unbinding kinetics of short and longer cTALEs are not due to differences in colocalization and FRET assays (Figure 3-figure supplement 1).

### A deterministic approach to modeling cTALE-DNA binding kinetics

To determine how the kinetic changes above are partitioned into underlying kinetic steps in binding, we fitted various kinetic models to the cumulative distributions for binding and unbinding. In addition to providing information about the mechanism of binding, this approach allows us to estimate the underlying microscopic rate constants and compare them for different constructs. This approach is generally applicable to studies of complex single molecule kinetics. Numerical integration was used to calculate the relative population of cTALE states as a function of time (Figures 5A-C and 5G-H), given a binding mechanism, an associated set of rate laws, and a set of initial conditions. Cumulative distributions of unbound dwell times represent the distribution of times single molecules spent in the unbound state before transitioning into the bound state, allowing us to split the kinetic scheme when fitting to single-molecule dwell times.

**Figure 5.**
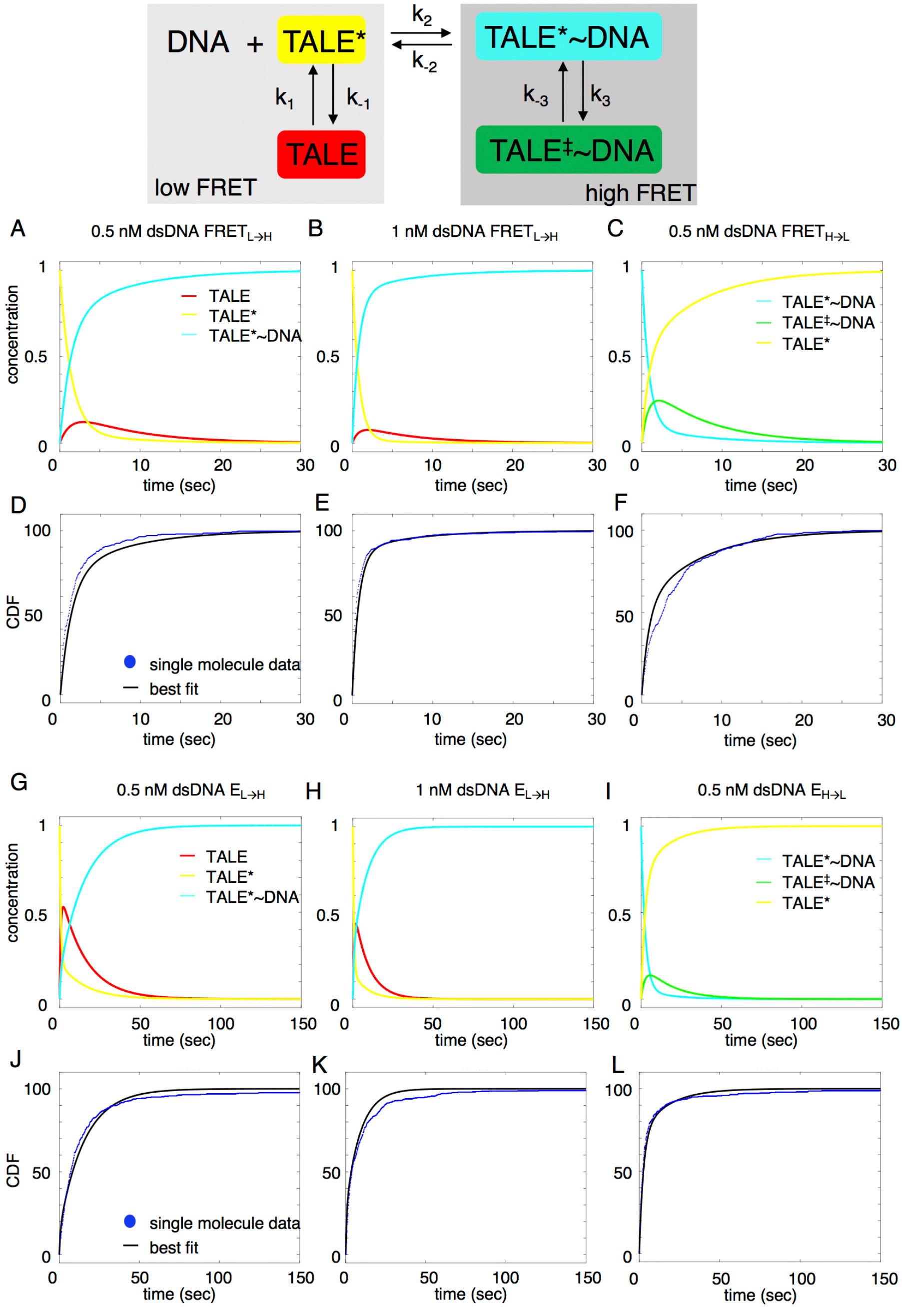
Deterministic simulations provide evidence for conformational heterogeneity in the unbound state. The model most consistent with data is shown at the top. Unbound TALEs can exist in DNA-binding competent (TALE*) or DNA-binding incompetent (TALE) states. DNA-bound TALEs can exist in short-lived (TALE*~DNA) or long-lived (TALE^‡^~DNA) DNA-bound states. Cumulative distributions of dwell-times (shown as blue points) from 8 repeat single-molecule time trajectories (A-F) and 16 repeat single-molecule time trajectories (G-L) were analyzed with the model (best-fit shown in black). (A-C and G-I) Populations of states as a function of time, generated by numerical integration in Matlab. (D-F and J-L) Cumulative distributions in blue circles and best fit lines are shown in black. Best-fit microscopic rate constants and 68% confidence intervals are listed in Table 1.

Among the various models tested, the model that is most consistent with the data has two unbound DNA-free states and two DNA-bound states. This is consistent with alternating laser experiments showing that DNA is only colocalized when cTALEs are in the high FRET state (Figure 3 - figure supplement 1). This four-state model includes a TALE isomerization step in the absence of DNA from a DNA-binding incompetent conformation (which we refer to as TALE) to DNA-binding competent conformation (which we refer to as TALE*). The DNA-binding competent TALE* conformer binds and unbinds DNA (called TALE* when DNA free and TALE*~DNA when DNA-bound).

Before unbinding, a fraction of TALE*~DNA isomerizes to a longer-lived DNA-bound state called TALE^‡^~DNA.

Based on this mechanism, the rate laws for binding are given in equations 1a-1d

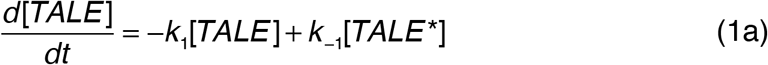

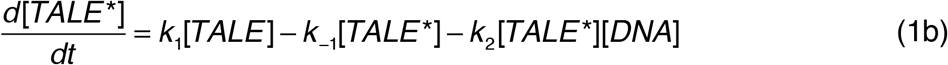

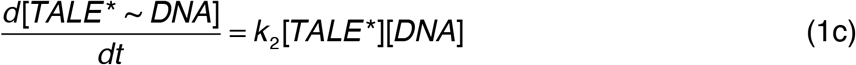

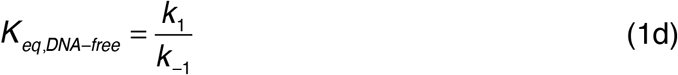

Since the single-molecule dwell-time histograms of the unbound states are insensitive to the isomerization after DNA binding, the equation describing the time evolution of the long-lived bound state (TALE^‡^~DNA) is not relevant to our analysis of unbound-state lifetimes.

To determine microscopic rate constants k_1_, k_-1_, and k_2_, equations 1a-1c were numerically integrated in Matlab, and the fraction of TALE*~DNA as a function of time was fitted to the low-FRET cumulative distributions (NcTALE_8_; Figure 5D-E) or to the no colocalization cumulative distributions (NcTALE_16_; Figure 5J-K). Microscopic rate constants were adjusted to reduce sum of the squared residuals between the concentration of TALE*~DNA (the direct product of binding) as a function of time and single-molecule cumulative distributions. In both cases, cumulative distributions at different bulk DNA concentrations were fitted globally. Initial fractions of TALE and TALE*~DNA were set to zero, and the initial fraction of TALE* was set to one. Confidence intervals (CI) were estimated by bootstrapping (Table 1; mean and 68% CI from 2000 or 8000 bootstrap iterations).

**Table 1.** Kinetic parameters obtained from deterministic simulation fits.

| | $k_1$ (sec<sup>-1</sup>) | $k_{-1}$ (sec<sup>-1</sup>) | $K_{eq, \text{DNA-free}}$ | $k_2$(sec<sup>-1</sup>nM<sup>-1</sup>) | $k_{-2}$ (sec<sup>-1</sup>) | $k_3$ (sec<sup>-1</sup>) | $k_{-3}$ (sec<sup>-1</sup>) | $K_{eq, \text{DNA-bound}}$<br>(nM<sup>-1</sup>) |
| --- | --- | --- | --- | --- | --- | --- | --- | --- |
| **NcTALE<sub>8</sub><sup>a</sup>** | 0.17<br>[0.16, 0.18] | 0.13<br>[0.12, 0.14] | 1.32<br>[1.26, 1.39] | 1.1<br>[1.08, 1.12] | 0.66<br>[0.65, 0.67] | 0.36<br>[0.35, 0.37] | 0.222<br>[0.218, 0.227] | 1.62<br>[1.58, 1.66] |
| **NcTALE<sub>12</sub><sup>b</sup>** | 0.135<br>[0.133, 0.137] | 1.26<br>[1.09, 1.34] | 0.11<br>[0.10, 0.12] | 0.31<br>[0.28, 0.33] | 0.130<br>[0.129, 0.131] | 0.043<br>[0.042, 0.044] | 0.0435<br>[0.0428, 0.0442] | 0.99<br>[0.97, 1.00] |
| **NcTALE<sub>16</sub><sup>a</sup>** | 0.26<br>[0.25, 0.27] | 0.43<br>[0.39, 0.47] | 0.61<br>[0.57, 0.64] | 0.39<br>[0.38, 0.41] | 0.299<br>[0.298, 0.300] | 0.074<br>[0.073, 0.076] | 0.078<br>[0.076, 0.079] | 0.96<br>[0.95, 0.97] |

68% confidence intervals shown in brackets are from 2,000<sup>a</sup> and 8,000<sup>b</sup> iterations of bootstrap analysis.

Rate laws for dissociation are given in equations 2a - 2d

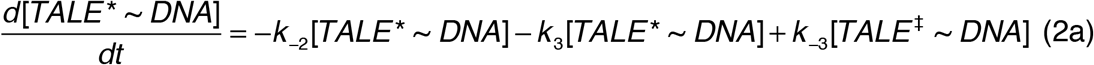

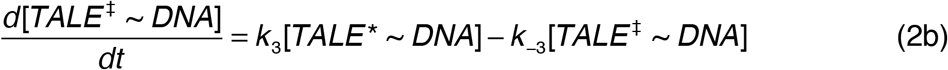

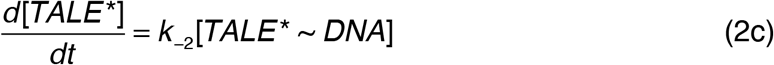

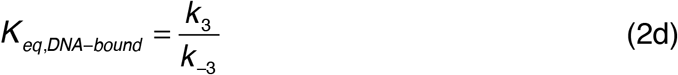

As with the system of equations above (1a-d), the equation describing the time evolution of the binding-incompetent free state (TALE) is not relevant to our analysis of bound-state lifetimes.

To determine microscopic rate constants k_-2_, k_-3_, and k_3_, equations 2a-2c were numerically integrated in Matlab, and the fraction of TALE* as a function of time was fitted to the high-FRET cumulative distributions (NcTALE_8_; Figure 5F) or to the low colocalization cumulative distributions (NcTALE_16_; Figure 5L). Microscopic rate constants were adjusted to reduce sum of the squared residuals between the concentration of TALE* (the direct product of dissociation) as a function of time and single-molecule cumulative distributions. In both cases, cumulative distributions at different bulk DNA concentrations were fitted globally. The initial fraction of TALE*~DNA conformer was set at one; all other initial fractions were set to zero. Confidence intervals were estimated by bootstrapping (Table 1; mean and 68% CI from 2000 iterations).

Fitted curves reproduce the experimental cumulative distributions for binding and unbinding (Figure 5), both for the short and long cTALE arrays, with reasonably small residuals, over a range of DNA concentrations. Generally, fitted rate constants have confidence intervals of 10% or smaller (Table 1).

Comparison of microscopic rate constants for 8, 12, and 16 repeats show some significant differences. The bimolecular microscopic binding rate constant, k_2_, is slightly larger for 8 repeats than for 12 and 16 repeats (1.1, 0.31, and 0.39 nM^−1^s^−1^for 8, 12, and 16 repeats respectively). However, microscopic unbinding rate constant, k-_2_, is higher for 8 repeat cTALEs (0.66 s^−1^for NcTALE_8_ versus 0.13 s^−1^for NcTALE_12_ and 0.299 s^−1^for NcTALE_16_). Also, bound state isomerization (interconversion between TALE*~DNA and TALE^‡^~DNA) is 5-10 times slower for 16 and 12 repeat cTALEs than 8 repeat cTALEs. The value of K_eq, DNA-free,_ which is a measure of the equilibrium proportion of the unbound TALE that is DNA-binding competent (TALE*) to that which is binding-incompetent (TALE), is larger for cTALEs with 8 repeats (K_eq, DNA-free_ = 1.32) than for cTALEs with 12 and 16 repeats (K_eq, DNA-free_ = 0.11 and K_eq, DNA-free_ = 0.61, respectively).

## Discussion

By measuring DNA-binding kinetics of cTALE arrays that form 0.7, 1, and 1.4 superhelical turns, we probe the functional relevance of locally unfolded TALE states. We describe a novel method to glean mechanistic details from complex single molecule kinetics. In our simplified cTALE system, we find conformational heterogeneity in both DNA free and DNA-bound states. We find that association is slowed in arrays containing one full turn of repeats or more. Because most natural and designed TALEs contain more than a full turn of repeats, this finding has important implications for design of high affinity TALE endonucleases (TALEN) molecules, suggesting that placement of destabilized repeats at specific positions may increase activity.

### cTALEs containing NS RVD bind DNA with high affinity

NS is an uncommon RVD in natural TALEs. Previous reports suggest that NS is fairly nonspecific, but may bind with higher affinity than other common RVDs (NG, NI, NN, and HD)(Miller et al., 2015). Our fitted rate constants can be used to calculate the apparent K_d_ (K_app_) calculated from using equation as follows:

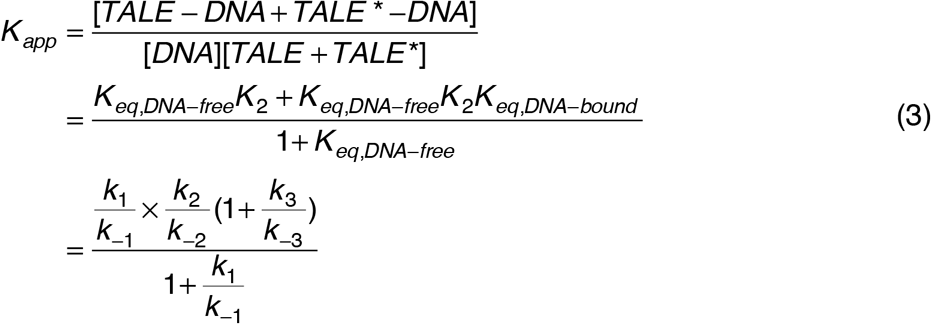

where *K*_2_ = *k*_2_ *k*_−2_.

Using fitted rate constants from Table 1 in the final equality in equation 3 gives values for K_app_ of 2.5 nM for the 8 repeat cTALE array, 0.5 nM for the 12 repeat cTALE array, and 1.0 nM for the 16 repeat cTALE array. Increasing the number of repeats has a modest affect on the apparent K_d_ due to the increased population of binding incompetent DNA free TALE in the 12, and 16 repeat arrays. This affinity change is small compared to a previous report studying length dependence on affinity of designed TALEs (dTALEs) showing the K_d_ of a dTALE decreased by a factor of two with the addition of only 1.5 repeats (Rinaldi et al., 2017).

TALEs are believed to read out sequence information from one strand (Boch et al., 2009). Due to the asymmetry of our DNA sequences (poly-dA base-paired with poly-dT), in principle, the FRET efficiency contains information on the binding orientation (and thus strand preference). However, based on the crystal structure of the DNA-bound state of TAL-effector PthXo1 (Mak et al., 2012), we estimate that the distance between the donor site of NcTALE_8_ (repeat 1) to the 5’ acceptor site on the DNA (Cy5-A_15_/T_15_) should be similar for both the dA-sense or dT-sense orientations (Figure 4-figure supplement 3A). Thus, the FRET data does not discriminate between the two modes of binding for the eight-repeat construct. However, for the 16 repeat NS RVD cTALE arrays, the PthXo1 model suggests very different distances (25 Å versus 73 Å for the dT-sense or dA-sense respectively, Figure 4-figure supplement 3B) between the donor site (TALE repeat 14) and the acceptor site (5’ Cy5-A_23_/T_23_). To restrict the number of binding positions available to longer cTALE arrays, the 23 base pair DNA used for NcTALE_16_ measurements (as well as DNA depicted in 4-figure supplement 3B) has the same number of additional base pairs as repeats (8 additional repeats and 8 additional base pairs) compared to the 15 base pair DNA used for NcTALE_8_ measurements (as well as DNA depicted in Figure 4-figure supplement 3A). While we limited the number of available binding positions, it may be possible for cTALEs to slide along DNA. However, taking into account the four repeat N-terminal capping domain, there are only three available base pairs in the bound complex. Thus we don’t expect the distance measurements to change by more than 10Å (~3 base pairs) if sliding occurs. The observation that there is colocalization but no measurable FRET when NcTALE_16_ is bound to DNA suggests that cTALEs containing the NS RVD prefer adenine (the dA-sense mode) compared with thymine bases, consistent with previous reports (Boch et al., 2009).

### Conformational heterogeneity in the unbound state may be caused by local unfolding

The cumulative distributions of dwell-times in Figure 3 provide clear evidence for conformational heterogeneity in both the free and DNA-bound cTALEs. Although the deterministic modeling supports such heterogeneity, puts it in the framework of a molecular model, and provides a means to determine the microscopic rate and equilibrium constants, such analysis provides little information about the structural nature of TALE conformational heterogeneity.

Figure 6 shows a model of cTALE conformational change consistent with DNA binding kinetics. In this model there are four TALE states. DNA-free cTALEs comprise both incompetent and binding competent states. DNA-bound cTALES comprise encounter and locked complexes with DNA. For 8 repeat cTALE arrays, the DNA-binding competent state is more highly populated than the DNA-binding incompetent state. In this reaction scheme, the DNA-binding incompetent state can be regarded as an off-pathway conformation that inhibits DNA binding (Figure 6A).

**Figure 6.**
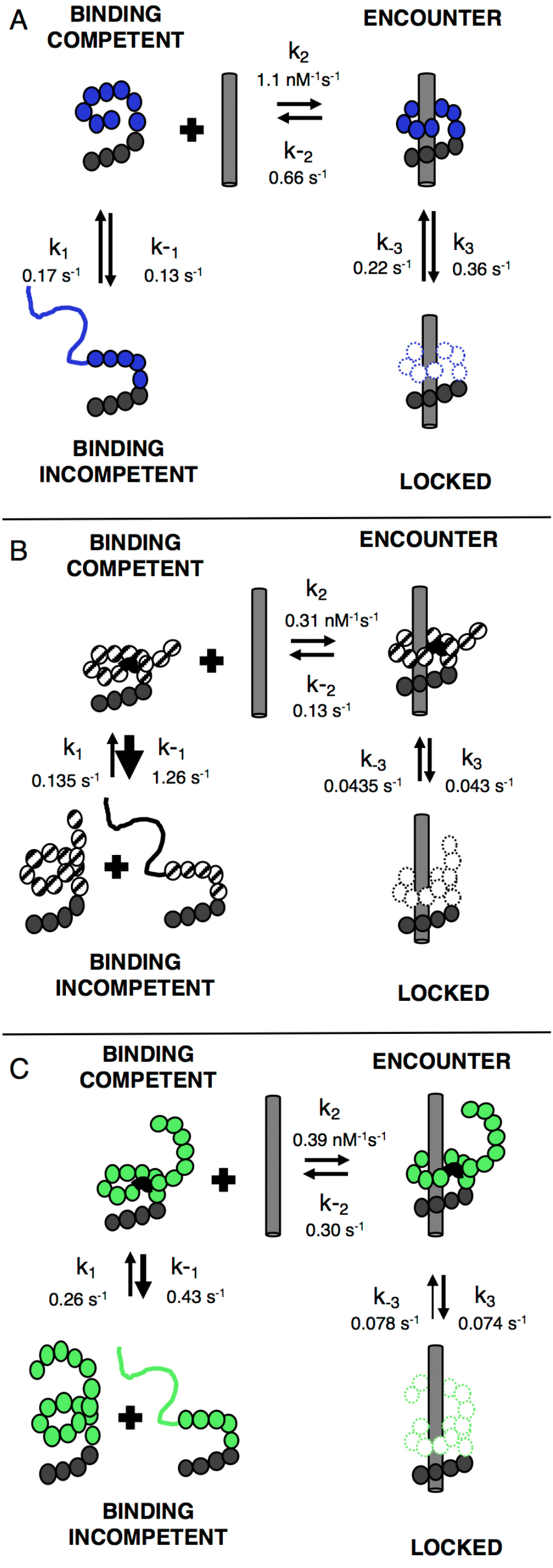
TALEs with multiple superhelical turns must break to bind DNA. Single-molecule FRET studies and deterministic modeling support a model where TALEs exist in four states: binding incompetent, binding competent, encounter complex, and locked. In this model, for TALEs that form less than one full superhelical turn (8 repeats, A), partly folded states are off-pathway and slow down binding. For longer TALEs that form one (12, B) or more (16, C) complete superhelical turns, partial unfolding is required for binding. DNA-bound TALEs form both encounter complexes and higher-affinity locked conformations. Dynamics of long (12 and 16-repeat; B-C) TALEs bound to DNA are significantly slower than for the shorter (8-repeat; A) TALE. **Figure supplement 1**. Urea and destabilizing mutations decrease apparent binding rate of cTALE_8_.

Because the 8 repeat cTALE array does not form multiple turns of a superhelix, unfolding to bind DNA is not required. In the model in Figure 6, the binding competent state is the fully folded conformation, whereas the binding incompetent state includes partly folded conformations. Consistent with this interpretation, increasing populations of partly folded states through addition of 1M urea and through entropy enhancing mutations decreases apparent binding rates of 8 repeat cTALEs (Figure 6-figure supplement 1). This is also consistent with a partly folded DNA-binding incompetent state in shorter cTALE arrays.

For 12 and 16 repeat cTALE arrays, the DNA-binding incompetent state is more highly populated than the DNA-binding competent state. In the model in Figure 6, the DNA-binding competent state is a high-energy conformation required for DNA binding (Figure 6B-C). Because 12 and 16 repeat cTALEs are expected to form 1 and 1.4 turns (excluding the N-terminal domain), we hypothesize that the binding competent state includes some partly folded states that allow access to DNA. Not all partly folded states open the array to access DNA; therefore, the binding incompetent state includes some nonproductive partly folded states in addition to the fully folded state.

In arrays containing 12 or more cTALEs, the binding competent and binding incompetent states likely include mixtures of many specific partly folded states. Because the types of partly folded states are unknown, connecting equilibria between binding competent and binding incompetent states to calculated partly folded equilibria (using folding free energies similar to Figure 1) is challenging. Future work towards understanding the structural characteristics of the binding competent state in TALE arrays of one or more turns would inform which partly folded states to include in the calculation, making this comparison meaningful. A better structural understanding of the DNA binding competent state will also allow an opportunity for precise placement of destabilized repeats in designed TALEN arrays which may enable more precise gene editing methodologies in both clinical and basic research applications.

### TALE functional instability presents a new mode of transcription factor binding

Here we demonstrate kinetic heterogeneity in DNA-bound and unbound TALE arrays, and we subsequently link the observed heterogeneity to partial unfolding of TALE arrays. We propose a model where binding requires partial unfolding of TALE arrays longer than one superhelical turn providing a functional role for previously observed moderate stability of TALE arrays. The functional instability described is particularly surprising given the small population of partly folded states which we expect to be DNA binding competent (partly folded states similar to internally unfolded and interfacially fractured states depicted in Figure 1A). Discovery of a functional role for the observed conformational heterogeneity is even more surprising, given the sequence identity of each of our repeats. Sequence heterogeneity in naturally occurring TALE arrays may further enable access to partly folded binding-competent states.

While it is well understood that many transcription factors sometimes undergo local folding transition upon DNA binding (Spolar and Record, 1994; Tsafou et al., 2018), the findings here indicate that for TALE arrays, the major conformer is fully folded, and must undergo a local unfolding transition in order to bind DNA. Taken together, these findings suggest a new mode of transcription factor binding and provide compelling evidence for functional instability in TALE arrays.

### Conformational heterogeneity in the bound state

Previous reports show that TALEs have multiple diffusional modes when searching nonspecific DNA (Cuculis et al., 2015). Our work suggests that cTALEs have multiple binding modes (encounter and locked states in Figure 6) indicating cTALEs undergo a conformational change also when bound to specific DNA sequences. Table 1 shows that microscopic rate constants for transition into and out of longer lived locked bound states become much slower in 12 and 16 repeat cTALEs compared with 8 repeat cTALEs (k_-3_ and k_3_). These rate constants decrease much more than the microscopic unbinding rate constant (the k_-2_ values are 0.66 s^−1^, 0.13 s^−1^, and 0.299 s^−1^for NcTALE_8_, NcTALE_12_, and NcTALE_16_ respectively) suggesting a large conformational change that depends on the number of repeats. Although the model does not provide information on structure of this conformational change, it is possible this conformational change involves a slinky motion to decrease helical rise. Consistent with this hypothesis, crystal structures of TALEs in the free and DNA-bound state show 11.5 repeats per turn in both states, but the helical rise decreases upon binding (Deng et al., 2012). Another possibility involves specific interaction with RVDs and bases in the major groove of DNA. Crystal structures show little deformation of DNA structure, so bending of DNA seems unlikely. While we can only hypothesize about the structural nature of the conformational changes, deterministic simulations show that cTALEs bind DNA through short encounters which occasionally become long-lived locked conformations (Figure 6). Taken together, these findings indicate that functional instability plays a crucial role in cTALE DNA binding, and demonstrate the importance of conformational dynamics in complex assembly.

## Materials and Methods

### Cloning, expression, purification, and labeling

Consensus TALE repeat constructs were cloned with C-terminal His_6_ tags via an in-house version of Golden Gate cloning (Cermak et al., 2011). TALE constructs were grown in BL21(T1R) cells at 37°C to an OD of 0.6-0.8 and induced with 1 mM IPTG. Following cell pelleting and lysis, proteins were purified by resuspending the insoluble material in 6M urea, 300 mM NaCl, 0.5 mM TCEP, and 10 mM NaPO_4_ pH 7.4. Constructs were loaded onto a Ni-NTA column. Protein was eluted using 250 mM imidazole and refolded during buffer exchange into 300 mM NaCl, 30% glycerol, 0.5 mM TCEP, and 10 mM NaPO_4_ pH 7.4.

Labelling of cTALE arrays followed a previously reported protocol (Rasnik et al., 2004). NcTALE_8_ and NcTALE_12_ were labeled at residue R30C in the first repeat, while NcTALE_16_ was labeled at residue R30C in the fourteenth repeat. 1 mg protein was loaded onto 500 uL NiNTA spin column. The column as washed with 10 column volumes of 300 mM NaCl, 0.5 mM TCEP, and 10 mM NaPO_4_ pH 7.4. Tenfold molar excess Cy3 maleimide dye was resuspended in 10 µL DMSO and added to column. The column was rocked at room temperature for 30 minutes, then at 4°C overnight. Cy3-labeled protein was eluted with 250 mM imidazole, 300 mM NaCl, 30% glycerol, 0.5 mM TCEP, and 10 mM NaPO_4_ pH 7.4. Protein was stored in 300 mM NaCl, 30% glycerol, 0.5 mM TCEP, and 10 mM NaPO_4_ pH 7.4 at −80°C.

### Oligonucleotides

Sequences used for binding studies were 5’-Cy5-A_15_-3’ and 5’ T_15_-3’ duplex (Cy5-A_15_/T_15_) for 8 repeat binding studies, and 5’-Cy5-A_23_-3’ and 5’ T_23_-3’ duplex (Cy5-A_23_/T_23_) for 12 and 16 repeat binding studies. DNA was annealed at 5 µM concentration with 1.2-fold molar excess unlabeled strand in 10 mM Tris pH 7.0, 30 mM NaCl.

### Single-molecule detection and data analysis

Biotinylated quarts slides and glass coverslips were prepared as previously described (Rasnik et al., 2004). Cy3-labeled cTALEs were immobilized on biotinylated slides taking advantage of neutravidin interaction with biotinylated α-penta•His antibody which binds the His_6_ cTALE tag. Slides were pretreated with blocking buffer (5 µL yeast tRNA, 5 µL BSA, 40 µL T50) before addition of 250 pM labeled cTALE. Cy5-labeled duplex DNA was mixed with imaging buffer (20 mM Tris pH 8.0, 200 mM KCl, 0.5 mg mL^−1^BSA, 1 mg mL^−1^glucose oxidase, 0.004 mg mL^−1^catalase, 0.8% dextrose and saturated Trolox ~1mg mL^−1^) and molecules were imagined using total internal reflection fluorescence microscopy. The time resolution was 50 msec for NcTALE_8_ and 100 msec for NcTALE_16_ and NcTALE_12_. Collection and analysis was performed as previously described (Roy et al., 2008).

### FRET histograms

A minimum of 20 short movies were collected, and the first 5 frames (50 msec exposure time) were used to generate smFRET histograms. FRET was calculated as I_A_/(I_A_+I_D_) where I_A_ and I_D_ are donor-leakage and background corrected fluorescence emission of acceptor (Cy5) and donor (Cy3) fluorophores. In competition experiments, unlabeled DNA with the same sequence as labeled DNA was mixed at indicated concentrations with labeled DNA prior to imaging.

### Dwell time analysis

Long movies were collected with 50 msec exposure time for NcTALE_8_ and 100 msec exposure time for NcTALE_16_ and NcTALE_16_. At least 20 representative traces at each DNA concentration were selected and dwell times were determined by fitting as previously described using HaMMy (McKinney et al., 2006) for FRET in NcTALE_8_. Dwell times in NcTALE_12_ and NcTALE_16_ colocalization experiments are determined by using a thresholding procedure for Cy5 excitation (Figure 4-figure supplement 1). The algorithm used to identify low and high emission states here is slightly different than previously described thresholding algorithms (Blanco and Walter, 2010). To reduce the number of incorrectly identified transitions arising from increased background and noise at higher Cy5-labeled DNA concentrations, a thresholding algorithm with two limits was implemented (see Figure 4-figure supplement 1). All FRET and colocalization data are well described by models with two distinct states (0.0 FRET and ~0.45 FRET as well as low colocalization and high colocalization). Dwell times of the same state (low versus high FRET or low versus high colocalization) for all traces at a given DNA concentration are compiled, and cumulative distribution is generated with spacing equal to imaging exposure time.

To determine apparent rate constants using model-independent analysis, cumulative distributions were fitted with single and double exponential decays (Figure 3 and 4). Observed rates from exponential decay fits were plotted as a function of DNA concentration. Apparent rate constants were calculated as slope of DNA concentration-dependent observed rates or average of DNA concentration-independent observed rates.

### Deterministic modeling

Equations 1a-1c and 2a-2c were numerically integrated using the ODE15s and ODE45 solver in MATLAB. Microscopic rate constants were adjusted to minimize the sum of squared residuals between ODE-determined concentration of bound or free TALE and single molecule cumulative distributions using lsqnonlin in MATLAB. 68% confidence intervals were estimated by performing 2000 or 8000 bootstrap iterations in which residuals from the best fit of the model to the data were randomly re-sampled (with replacement) and re-fitted. All scripts and source data required to run this MATLAB program called <u>**De**</u>terminstic <u>**M**</u>odeling for <u>**A**</u>nalysis of complex <u>**S**</u>ingle molecule <u>**K**</u>inetics (DeMASK) are publicly available on GitHub at https://github.com/kgeigers/DeMASK.

## Acknowledgements

The authors thank members of the Barrick and Ha lab for their input on this work. The authors acknowledge the support of the Center for Molecular Biophysics at Johns Hopkins and Dr. Katherine Tripp for instrumental and technical support. Support to KGS was provided by NIH training grant T32-GM008403. Support for this project was provided by NIH grant 1R01-GM068462 to DB and GM112659 to TH.

## Supplemental Material

**Figure 3-figure supplement 1.**
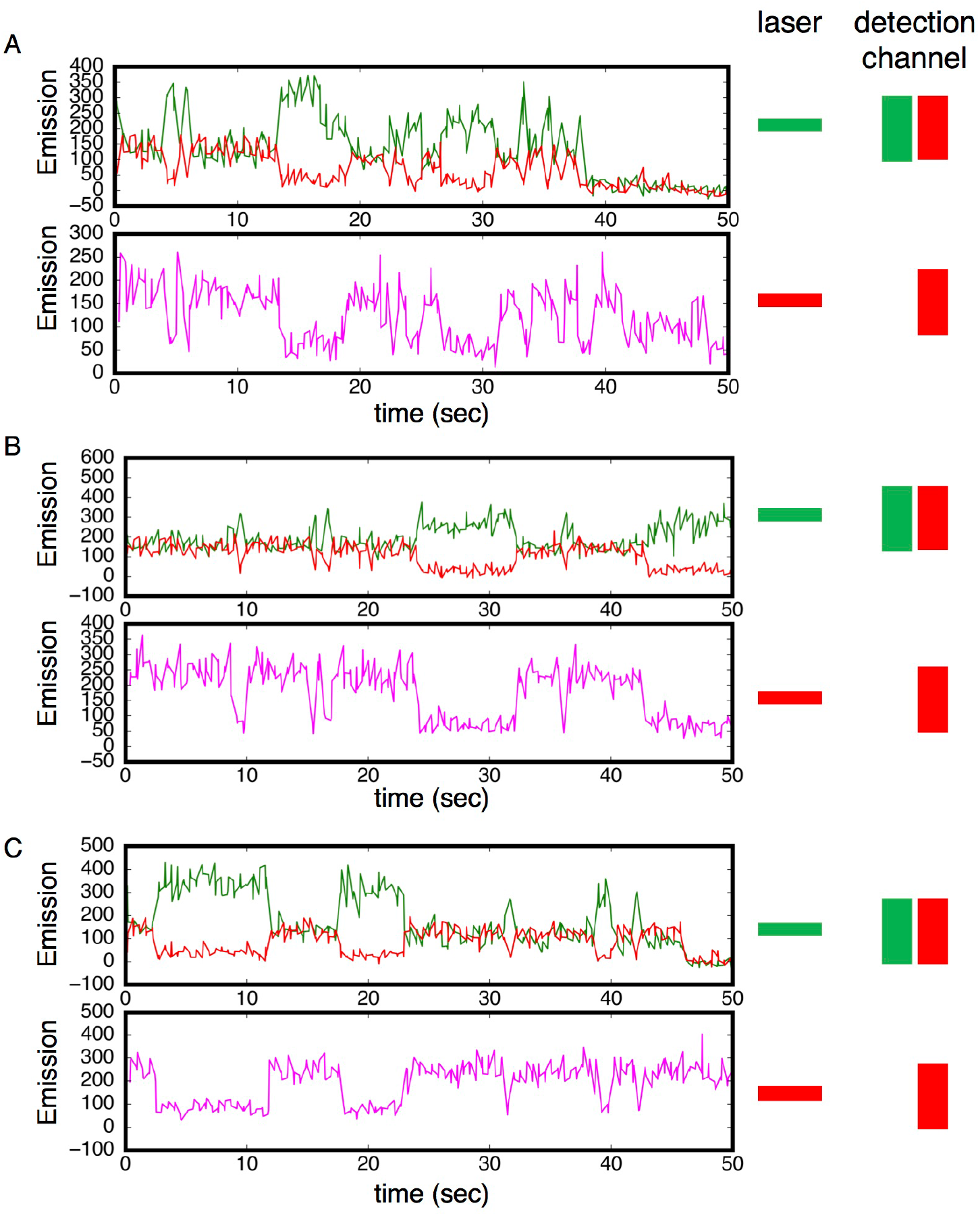
Alternating laser experiments show agreement between cTALE_8_ FRET and colocalization kinetics. (A-C) Three representative time trajectories. In these trajectories, excitation alternated between red and green with 250 msec at each color. Top panels show detection of red and green light resulting from green laser excitation. High red (and low green) fluorescence result from FRET. Lower panels show detection of red light resulting from red laser excitation. There is strong correlation between periods of high FRET in the upper panels (high red fluorescence, low green fluorescence) and periods of high red fluorescence (from DNA binding) in the lower panel. Conditions: 200 mM KCl, 20 mM Tris pH 8.0.

**Figure 3 – figure supplement 2.**
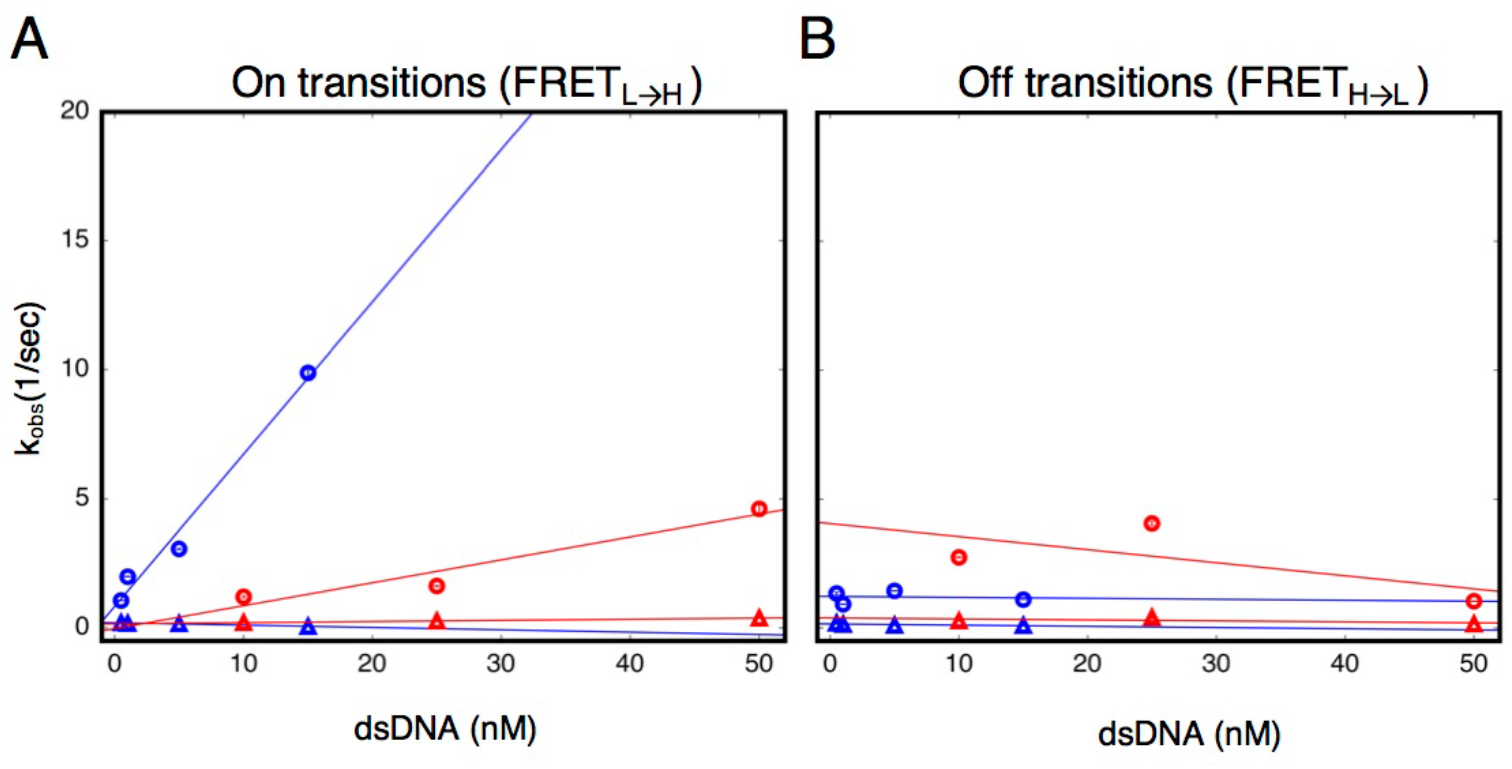
cTALEs do not slide onto ends of short dsDNA. Apparent association and dissociation rate constants as a function of DNA concentration for an 8 repeat cTALE array (NcTALE_8_) binding to uncapped (blue) and capped (red) DNA, measured by single molecule FRET dwell time analysis. (A) Rate constants for conversion from the low to the high FRET state (FRET_L→H_). Rate constants for the faster phase increase with DNA concentration (circles), indicating a bimolecular event. The DNA concentration-dependence is stronger (larger slope, k=0.59 ± 0.08 nM^−1^sec^−1^) with uncapped DNA than with capped DNA (k=0.09 ± 0.05 nM^−1^sec^−1^), which is likely a result of the faster diffusion of small, uncapped DNA (10 kDa) compared to large, capped DNA (320 kDa). Rate constants for the slower phase are not DNA-concentration dependent (triangles). (B) Rate constants for conversion from the high to the low FRET state (FRET_H→L_). Neither phase shows a DNA concentration dependent, indicating unimolecular steps. Conditions: 200 mM KCl, 20 mM Tris pH 8.0.

**Figure 4-figure supplement 1.**
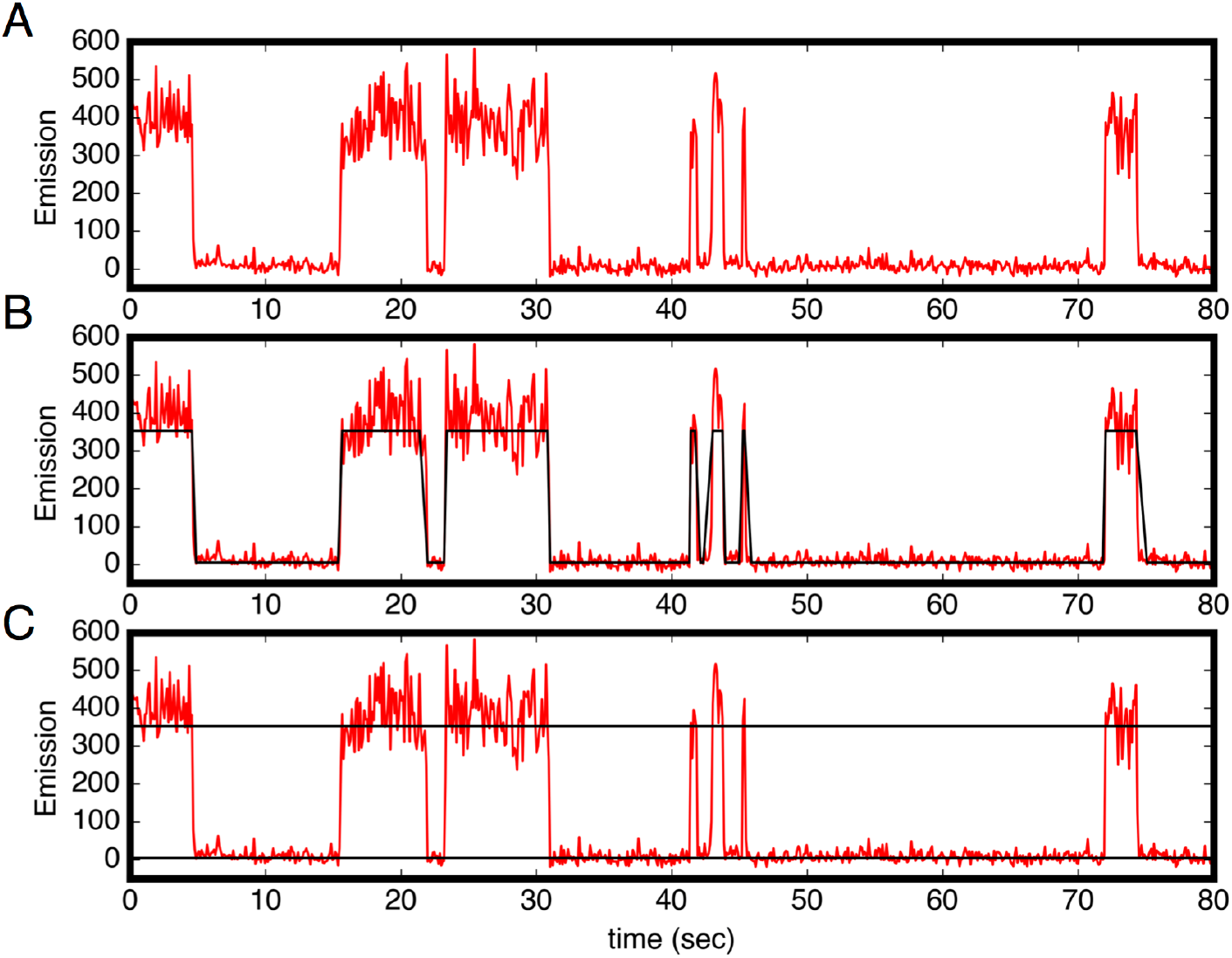
Colocalization trajectories show TALE-DNA binding and unbinding events. (A-C) One representative time trajectory and colocalization analysis for a sixteen repeat TALE, NcTALE_16_, incubated with 1 nM Cy5-A_15_/T_15_. In the colocalization protocol, both the green and red lasers were used in the first ten frames (not shown) to identify single molecule locations. For all subsequent frames, Cy5-labeled DNA was continuously excited with the red laser. Red light emission images (from Cy5) were collected with time steps of 100 msec. Trajectories were generated from sites of red/green colocalization in the first ten frames, and were corrected for background Cy5 emission (red trajectories above). For each trajectory, we set two thresholds for low and high Cy5 emission (C). A molecule is designated to be in the high emission (DNA-bound) state above the high threshold, and in the low emission state below the low threshold. From this state-assignment procedure, dwell-times were determined in the high (bound) and low (unbound) states (C). Conditions: 200 mM KCl, 20 mM Tris pH 8.0.

**Figure 4-figure supplement 2.**
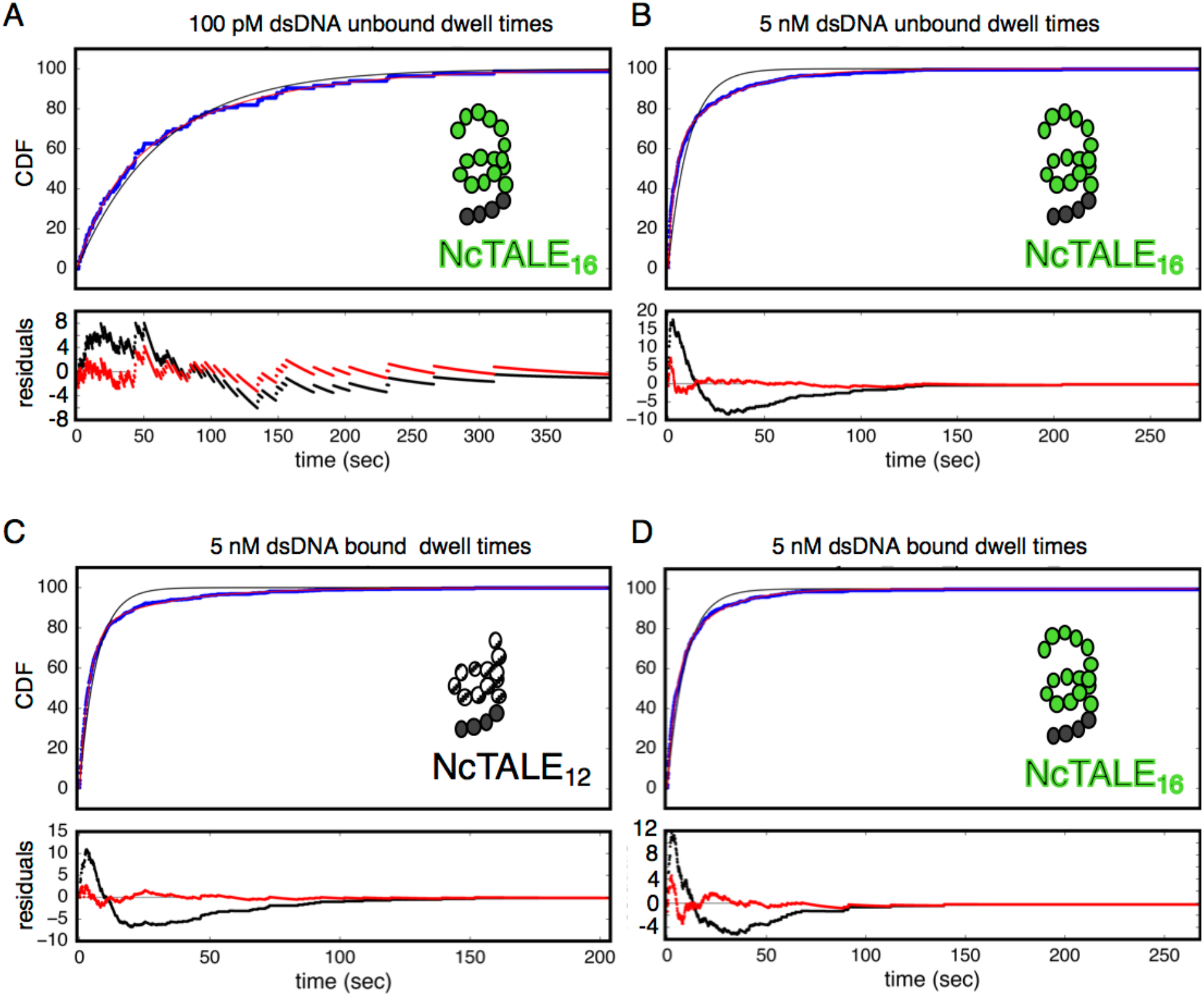
Bound and unbound lifetimes of 16- and 12-repeat TALE proteins are consistent with multiphasic binding and unbinding. (A, B) Cumulative distributions of low-FRET dwell times (blue circles) for a 16-repeat construct at low and high DNA concentration. Fits to single-exponentials (black) show large nonrandom residuals (lower panels) which are most pronounced at high DNA concentrations, consistent with the binding heterogeneity observed in eight repeat cTALEs (Figure 3). Double-exponentials (red) give smaller, more uniform residuals. (C, D) Cumulative distributions of high-FRET dwell times (blue circles) for 16- and 12-repeat constructs at high DNA concentration. Fits to single-exponentials (black) show large nonrandom residuals (lower panels), consistent with the dissociation heterogeneity observed in eight repeat cTALEs. Double-exponentials (red) give smaller, more uniform residuals.

**Figure 4-figure supplement 3.**
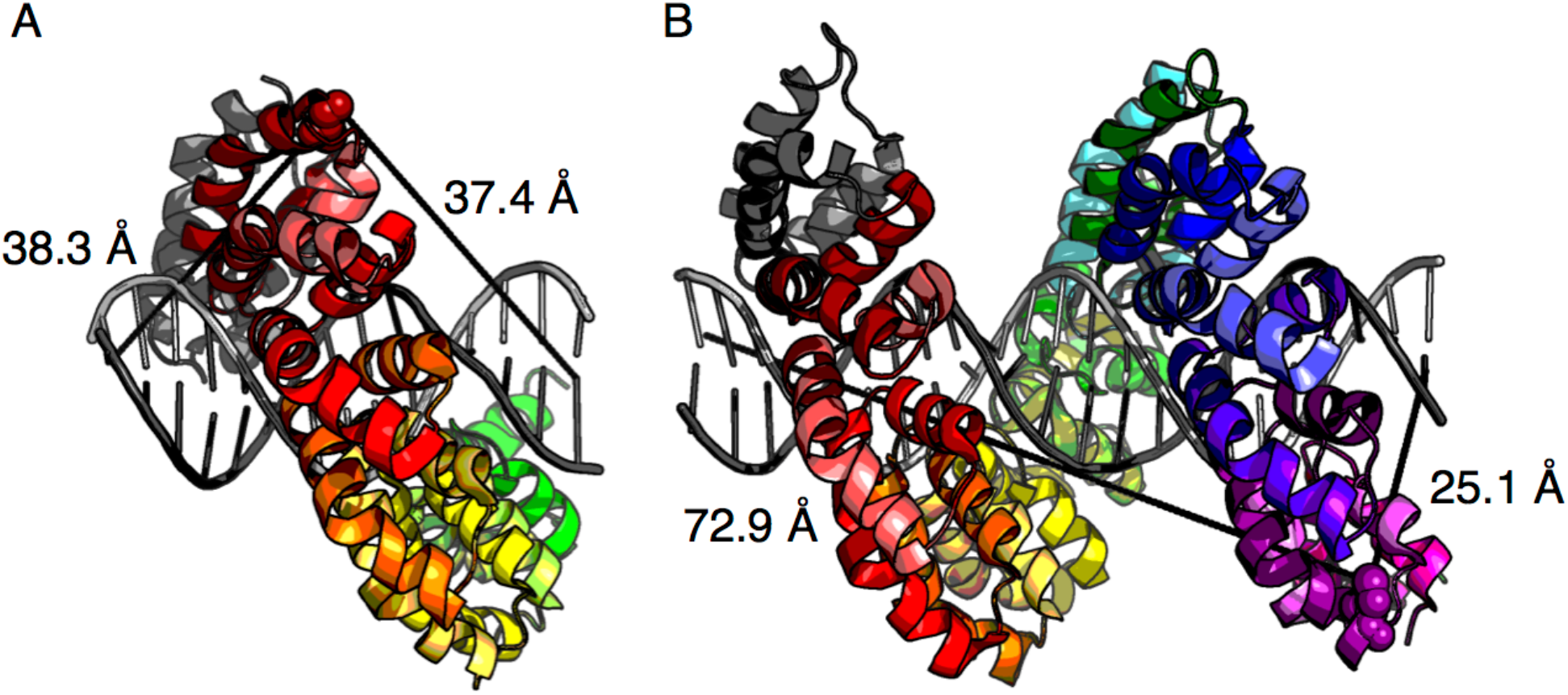
Distance estimates between labeling sites for NcTALE_8_ and NcTALE_16_ and the 5’ ends of bound DNA. The crystal structure of PthXo1 (PDB: 3UGM) (Mak et al., 2012) in complex with DNA (colored by repeat) as a model of donor-acceptor fluorophore distances. (A) The distance from C30 in the first repeat (the site of labelling in NcTALE_8_) to the 5’ base of the sense strand (light grey) is 38.3Å; the distance to the 5’ base of the antisense strand (dark grey) is 37.4Å. (B) The distance from C30 in the fourteenth repeat (the site of labelling in NcTALE_16_) to the 5’ base of the sense strand (light grey) is 72.9Å; the distance to the 5’ base of the antisense strand (dark grey) is 25.1Å. Note that although there is likely to be some variation these distances since the DNAs used here are longer (15 and 23 bases) than the TALE arrays (8 and 16 repeats, each with a four-repeat N-capping domain), the differences in donor-acceptor distances for the 16 repeat construct (B) are likely to be robust to registry shifts of a few bases.

**Figure 6-figure supplement 3 1.**
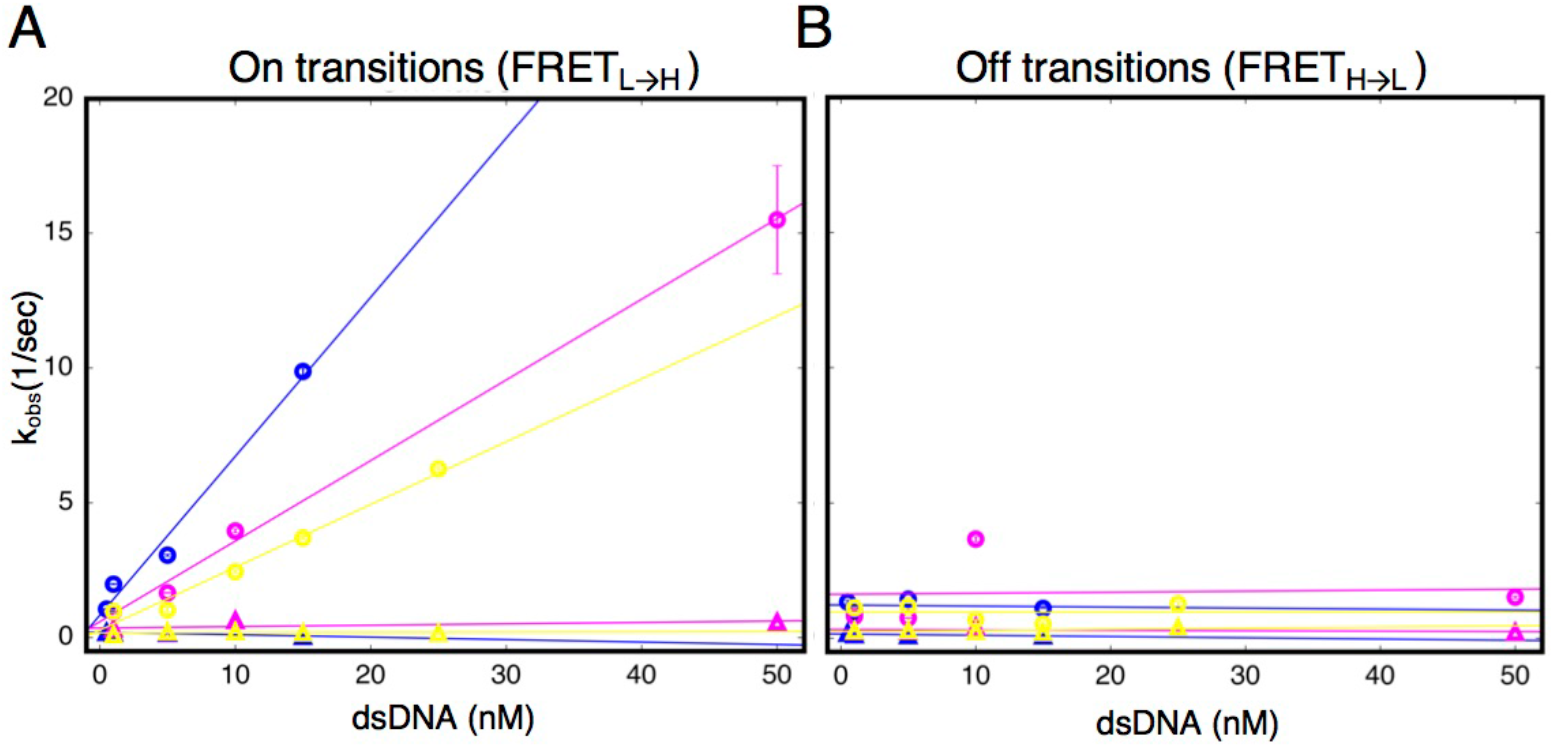
Urea and destabilizing mutations decrease apparent binding rate of cTALE_8_. (A) Apparent association rate constants as a function of DNA concentration for an 8 repeat cTALE in 0 M urea (blue), 1 M urea (pink), and with destabilizing point mutations (yellow; mutations substitute leucine at position 1 in the fourth repeat with a glycine). The DNA concentration dependence is strongest (steepest slope) in the absence of urea and destabilizing point substitutions (circles). Rates in the slower phase (triangles) appear unaffected by urea or mutational destabilization. (B) Apparent dissociation rate constants as a function of DNA concentration (phase 1 shown in circles, and phase 2 shown in triangles). Rates in the both phases appear unaffected by urea and mutational destabilization. Conditions: 200 mM KCl, 20 mM Tris pH 8.0.

## Source Data

**Figure 3-source data 1. List of values used to construct long time trajectories.** Long time trajectories showing transitions between low- and high-FRET states (efficiencies of 0 and 0.45) for 1 nM dsDNA (source data 1A) and 15 nM dsDNA (source data 1B) depicted in Figure 3A-B.

**Figure 3-source data 2. List of values used to construct CDFs.** Cumulative distributions of low-FRET (source data 2A) and high-FRET (source data 2B) dwell times depicted in Figure 3C-D.

